# Genetic analysis of SARS-CoV-2 isolates collected from Bangladesh: insights into the origin, mutation spectrum, and possible pathomechanism

**DOI:** 10.1101/2020.06.07.138800

**Authors:** Md Sorwer Alam Parvez, Mohammad Mahfujur Rahman, Md Niaz Morshed, Dolilur Rahman, Saeed Anwar, Mohammad Jakir Hosen

**Affiliations:** Department of Genetic Engineering & Biotechnology, Shahjalal University of Science & Technology, Sylhet 3114, Bangladesh; Department of Medical Genetics, Faculty of Medicine and Dentistry, University of Alberta, 8440 112 St. NW, Edmonton, AB T6G 2R7, Canada

**Keywords:** COVID-19, SARS-CoV-2, Bangladeshi isolates, genome, spike protein, mutation, ACE2 receptor

## Abstract

As the coronavirus disease 2019 (COVID-19), caused by the severe acute respiratory syndrome coronavirus-2 (SARS-CoV-2), rages across the world, killing hundreds of thousands and infecting millions, researchers are racing against time to elucidate the viral genome. Some Bangladeshi institutes are also in this race, sequenced a few isolates of the virus collected from Bangladesh. Here, we present a genomic analysis of 14 isolates. The analysis revealed that SARS-CoV-2 isolates sequenced from Dhaka and Chittagong were the lineage of Europe and the Middle East, respectively. Our analysis identified a total of 42 mutations, including three large deletions, half of which were synonymous. Most of the missense mutations in Bangladeshi isolates found to have weak effects on the pathogenesis. Some mutations may lead the virus to be less pathogenic than the other countries. Molecular docking analysis to evaluate the effect of the mutations on the interaction between the viral spike proteins and the human ACE2 receptor, though no significant interaction was observed. This study provides some preliminary insights into the origin of Bangladeshi SARS-CoV-2 isolates, mutation spectrum and its possible pathomechanism, which may give an essential clue for designing therapeutics and management of COVID-19 in Bangladesh.

## Introduction

The coronavirus disease 2019 (COVID-19) is an infectious disease caused by severe acute respiratory syndrome coronavirus 2 (SARS-CoV-2). Common symptoms of the disease include fever, cough, fatigue, shortness of breath, nausea, vomiting, and diarrhea. The disease has emerged as a critical, rapidly evolving global health crisis [1-3]. More than 6.5 million people have contracted the virus, and nearly 400 thousand have died [4, 5]. In Bangladesh, the COVID-19 was first reported on 7 March by the Institute of Epidemiology Disease Control and Research (IEDCR) [6]. Until the end of March, the infection rate was sort of low; however, as the non-therapeutic prevention measures enforced by the government faced enormous challenges, the infection rate raised drastically in April and kept on rising [7]. The people did not maintain the social distancing enforced by the government and trend to gather in crowded places [8]. Moreover, an inadequacy of testing for COVID-19 diagnosis is a common criticism in Bangladesh [9]. As of 5 June 2020, nearly 65 thousand confirmed cases were reported, with a total of 846 deaths in Bangladesh [10].

SARS-CoV-2 is a positive-stranded RNA virus with a genome of ∼ 30kb, encodes structural and non-structural proteins. Like other RNA viruses, the SARS-CoV-2 is prone to frequent mutations, which makes it challenging to develop therapeutics and vaccines against the virus [11, 12]. Sequence information of both the pathogen and the host would greatly facilitate an effective therapeutic strategy or vaccine development [13]. Analysis of the genome sequences obtained from a vast array of isolates collected from different regions could provide an idea about the efficacy of the vaccines being developed [14]. Henceforth, researchers across the world are running against time to unravel the genomic insights into the virus.

Till 2 June 2020, some thirty-five thousand genome sequences of SARS-CoV-2 has been submitted from different countries, where most of the sequence have come from European countries (∼20000). About 7000 complete genome sequences have been submitted from the USA while China has submitted ∼850 genome sequences. In Bangladesh, 16 isolates of the virus have been sequenced and deposited in GSAID (Global Initiative on Sharing Avian Influenza Data) database till 20^th^ May 2020. Unfortunately, there is yet a study on the genomics of the SARS-CoV-2 in Bangladeshi isolates.

This study aimed to provide some preliminary insights into the genetic structure of all isolates reported in Bangladesh along with the mutational spectrum. It presents the first study on SARS-CoV-2 genomes obtained from Bangladesh, which, in broader terms, would help the therapeutic strategy development and vaccination programs against the virus in the country.

## Materials and Methods

### Retrieval of the SARS-CoV-2 Genome Sequences

Till 20^th^ May, genome sequence of 16 Bangladeshi SARS-CoV-2 isolates were found deposited in the GSAID, however genome sequence of 2 isolates were found incomplete. Thus, all 14 complete genome sequences of the reported isolates of SARS-CoV-2 in Bangladesh were retrieved from the GISAID database (https://www.gisaid.org/). As many of the Bangladeshi people return during the COVID-19 outbreak mainly from China, India, Saudi Arabia, Spain, Italy, Japan, Qatar, Canada, Kuwait, USA, France, Sweden, and Switzerland, the first deposited genome sequence of those countries were also retrieved. Sequence information of the first isolate collected from China was considered as a reference for further analysis.

### Identification of Nucleotide Variations in Bangladeshi Strain

We performed multiple sequence alignment using Clustal Omega [15, 16], and the sequence of the strain China [EPI_ISL_402124] was used as a reference genome. The alignment file was analyzed using MVIEW program of Clustal Omega [17]. Only variations in the coding regions were analyzed in this study.

### Prediction of Viral Genome and Identification of Selected Genes

FGENESV of SoftBerry (http://linux1.softberry.com/berry.phtml), which is a Trained Pattern/Markov chain-based viral gene prediction tools, was adopted for the prediction of the genes as well as the proteins from the viral genomes. Each predicted protein (for each viral genomes) was identified using the Basic Local Alignment Search Tool (BLAST), at the interface of the National Center for Biotechnology Information (NCBI). The identity of each protein was evaluated compared to the proteins of the reference strain [18].

### Detection of Mutation Spectrum

Again, Clustal Omega was used for the multiple sequence alignment of each protein, which further analyzed by MVIEW. The amino acid variations were identified in each protein comparing to the protein of the reference strain. Further, both nucleotide variations and amino acid variations were compared to study the types of mutations.

### Prediction of Mutational Effects

The structural and functional effects of the missense variants, along with the stability change, were analyzed using different prediction tools. I-mutant was employed to analyze the stability change where all the parameters were kept in default [19]. Additionally, Mutpred2 was adopted to predict the molecular consequences and functional effect of these mutations [20].

### Homology Modeling of Spike Proteins and Validation

The BLASTp program at the NCBI interface (link) was used to find the most suitable template for homology modeling. Blasting against the protein databank reservoir (PDB) identified spike protein (Human) with PDB ID: 6VSB as a suitable template, as it has 99.59% sequence similarity and 94% coverage with the target sequence. The homology modeling of all mutant spike proteins along with the spike protein of the reference was done using SWISS-MODEL [21]. The validation of the predicted model was done by adopting Rampage and ERRAT [22, 23].

### Molecular Docking of Spike Protein with ACE2 Receptor

The molecular docking approach was employed to investigate the interaction of mutant spike protein with the human ACE2 receptor. First, the crystal structure of human ACE2 (PDB ID 6D0G) was obtained from Protein Data Bank, and PyMOL was used to clean the structure to remove all the complex molecules and water [24, 25]. The HDOCK webserver was used for prediction of the interaction between Spike protein and human ACE2 receptor through the protein-protein molecular docking [26]. PyMOL was also used for the visualization of docking interactions.

## Results

### Retrieved Genome Sequence of the SARS-CoV-2

A total number of 14 complete genome sequences of the SARS-CoV-2 isolates from Bangladesh and 12 genome sequence from the isolates of other countries (China, India, Saudi Arabia, Spain, Italy, Japan, Qatar, Canada, Kuwait, USA, France, Sweden, and Switzerland) have been retrieved from GSAID. The strain of Wuhan accession number with EPI_ISL_402124 was considered as the reference strain.

### Phylogenetic Tree Analysis

Phylogenetic tree analysis revealed that all the selected Bangladeshi isolates could be divided into two main groups, where one group shared a common ancestor with Saudi Arabia (Fig 1). The other group found to have a similarity with the strain from Switzerland, and it could be subdivided into two groups. In one subdivision, four isolates clustered with the strain from Spain, while the other group consisted only of the three Bangladeshi isolates. All the Bangladeshi isolates centered and shared a common ancestor with India and the USA.

**Fig 1:**
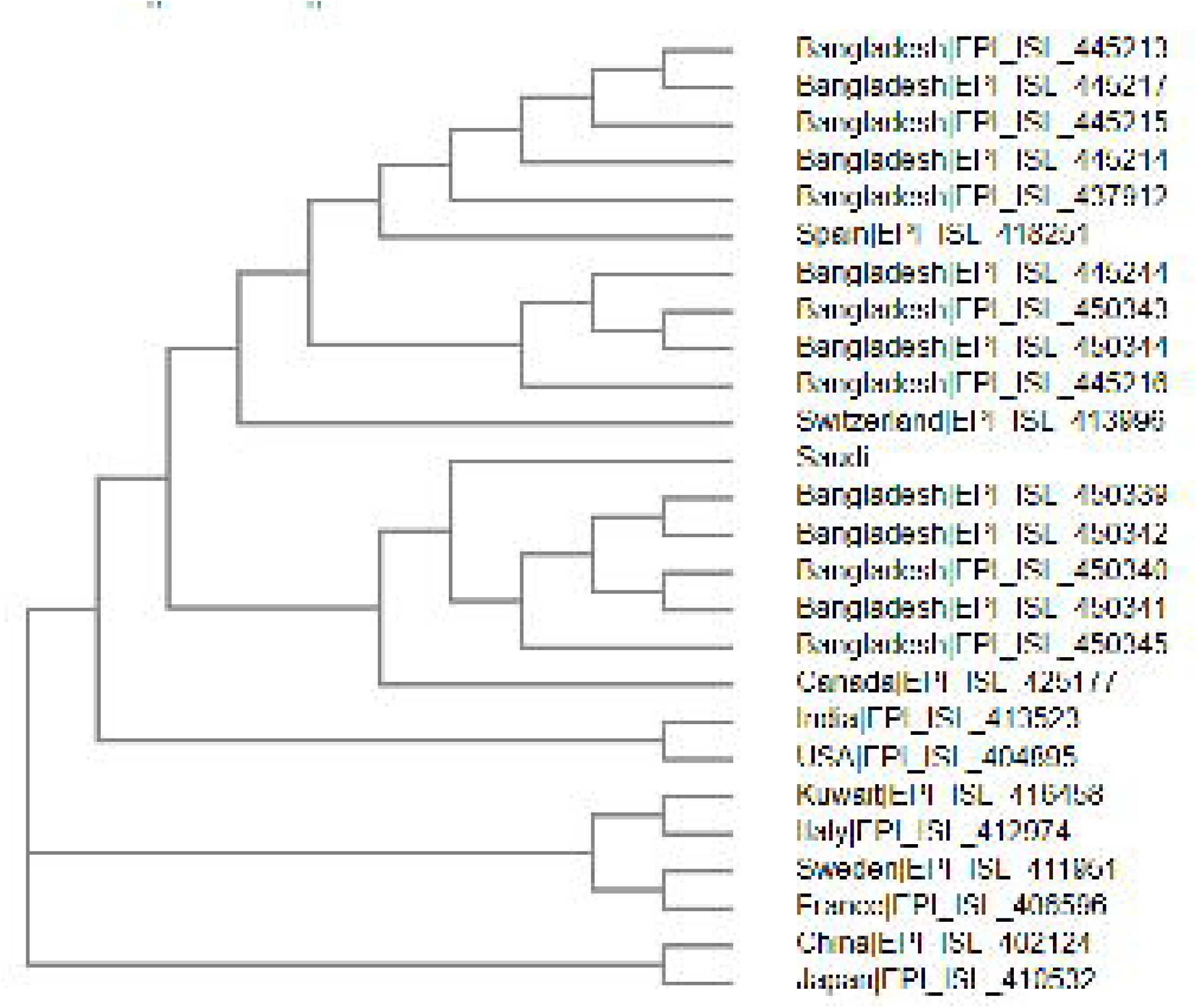
Phylogenetic tree of the 14 Bangladeshi isolates of the virus along with other 12 countries

### Predictions of the Genes and Proteins

FGENESV predicted the presence of 12 genes in the reference. Interestingly, all except five isolates (EPI_ISL_445213, EPI_ISL_445214, EPI_ISL_450342, EPI_ISL_450343, and EPI_ISL_450344) of Bangladesh also showed a similar result. Both isolates EPI_ISL_445213 and EPI_ISL_445217 found to have ten genes (missing of ORF7a and ORF10 genes) and isolate EPI_ISL_450343 and EPI_ISL_450344 have 11 genes (missing ORF8 gene). Multiple sequence alignment revealed that most of the variation in Bangladeshi isolates occurred in the ORF1a polyprotein, surface glycoproteins, and nucleocapsid phosphoprotein. Remarkably, envelope glycoprotein, ORF6, ORF8, and ORF10 were found 100% identical in most of the isolates compared to the reference sequence (Table 1).

**Table 1:**
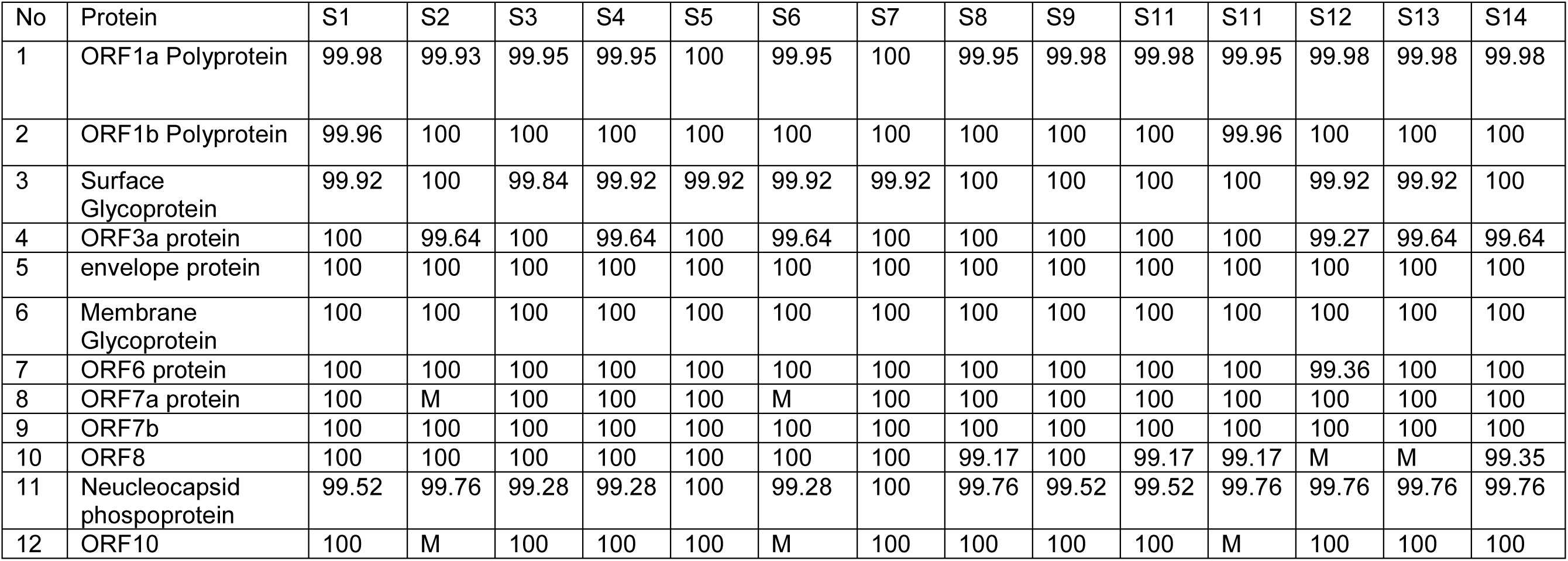
Predicted number of genes and identity compared to the reference strain. (Legends: S1: EPI_ISL_437912; S2: EPI_ISL_445213; S3: EPI_ISL_445214; S4: EPI_ISL_445215; S5: EPI_ISL_445216; S6: EPI_ISL_445217; S7: EPI_ISL_445244; S8: EPI_ISL_450339; S9: EPI_ISL_450340; S10: EPI_ISL_4503441; S11: EPI_ISL_450342; S12: EPI_ISL_450343; S13: EPI_ISL_450344; S14: EPI_ISL_450345; M: Missing)

| No | Protein | S1 | S2 | S3 | S4 | S5 | S6 | S7 | S8 | S9 | S11 | S11 | S12 | S13 | S14 |
| --- | --- | --- | --- | --- | --- | --- | --- | --- | --- | --- | --- | --- | --- | --- | --- |
| 1 | ORF1a Polyprotein | 99.98 | 99.93 | 99.95 | 99.95 | 100 | 99.95 | 100 | 99.95 | 99.98 | 99.98 | 99.95 | 99.98 | 99.98 | 99.98 |
| 2 | ORF1b Polyprotein | 99.96 | 100 | 100 | 100 | 100 | 100 | 100 | 100 | 100 | 100 | 99.96 | 100 | 100 | 100 |
| 3 | Surface Glycoprotein | 99.92 | 100 | 99.84 | 99.92 | 99.92 | 99.92 | 99.92 | 100 | 100 | 100 | 100 | 99.92 | 99.92 | 100 |
| 4 | ORF3a protein | 100 | 99.64 | 100 | 99.64 | 100 | 99.64 | 100 | 100 | 100 | 100 | 100 | 99.27 | 99.64 | 99.64 |
| 5 | envelope protein | 100 | 100 | 100 | 100 | 100 | 100 | 100 | 100 | 100 | 100 | 100 | 100 | 100 | 100 |
| 6 | Membrane Glycoprotein | 100 | 100 | 100 | 100 | 100 | 100 | 100 | 100 | 100 | 100 | 100 | 100 | 100 | 100 |
| 7 | ORF6 protein | 100 | 100 | 100 | 100 | 100 | 100 | 100 | 100 | 100 | 100 | 100 | 99.36 | 100 | 100 |
| 8 | ORF7a protein | 100 | M | 100 | 100 | 100 | M | 100 | 100 | 100 | 100 | 100 | 100 | 100 | 100 |
| 9 | ORF7b | 100 | 100 | 100 | 100 | 100 | 100 | 100 | 100 | 100 | 100 | 100 | 100 | 100 | 100 |
| 10 | ORF8 | 100 | 100 | 100 | 100 | 100 | 100 | 100 | 99.17 | 100 | 99.17 | 99.17 | M | M | 99.35 |
| 11 | Neucleocapsid phosphoprotein | 99.52 | 99.76 | 99.28 | 99.28 | 100 | 99.28 | 100 | 99.76 | 99.52 | 99.52 | 99.76 | 99.76 | 99.76 | 99.76 |
| 12 | ORF10 | 100 | M | 100 | 100 | 100 | M | 100 | 100 | 100 | 100 | M | 100 | 100 | 100 |

### Mutation Spectrum of Bangladeshi SARS-CoV-2 isolates

Analysis of all 14 Bangladeshi isolates revealed a total of 42 single nucleotide variants (Fig 2); 24 of them were nonsynonymous missense in character. Besides, three large deletions were also found in those isolates (Table 2). Among the deletions, two deletions were responsible for the deletion of ORF7a in EPI_ISL_445213 and EPI_ISL_445217 isolates. Another large deletion from nucleotide 27911 to 28254, occurred in EPI_ISL_450343 and EPI_ISL_450344 isolates, responsible for the deletion of ORF8 in both isolates. Surprisingly, three consecutive mutations were found at nucleotide position 28882 to 28884; resulted in two amino acids substitution in nucleocapsid phosphoprotein.

**Table 2:**
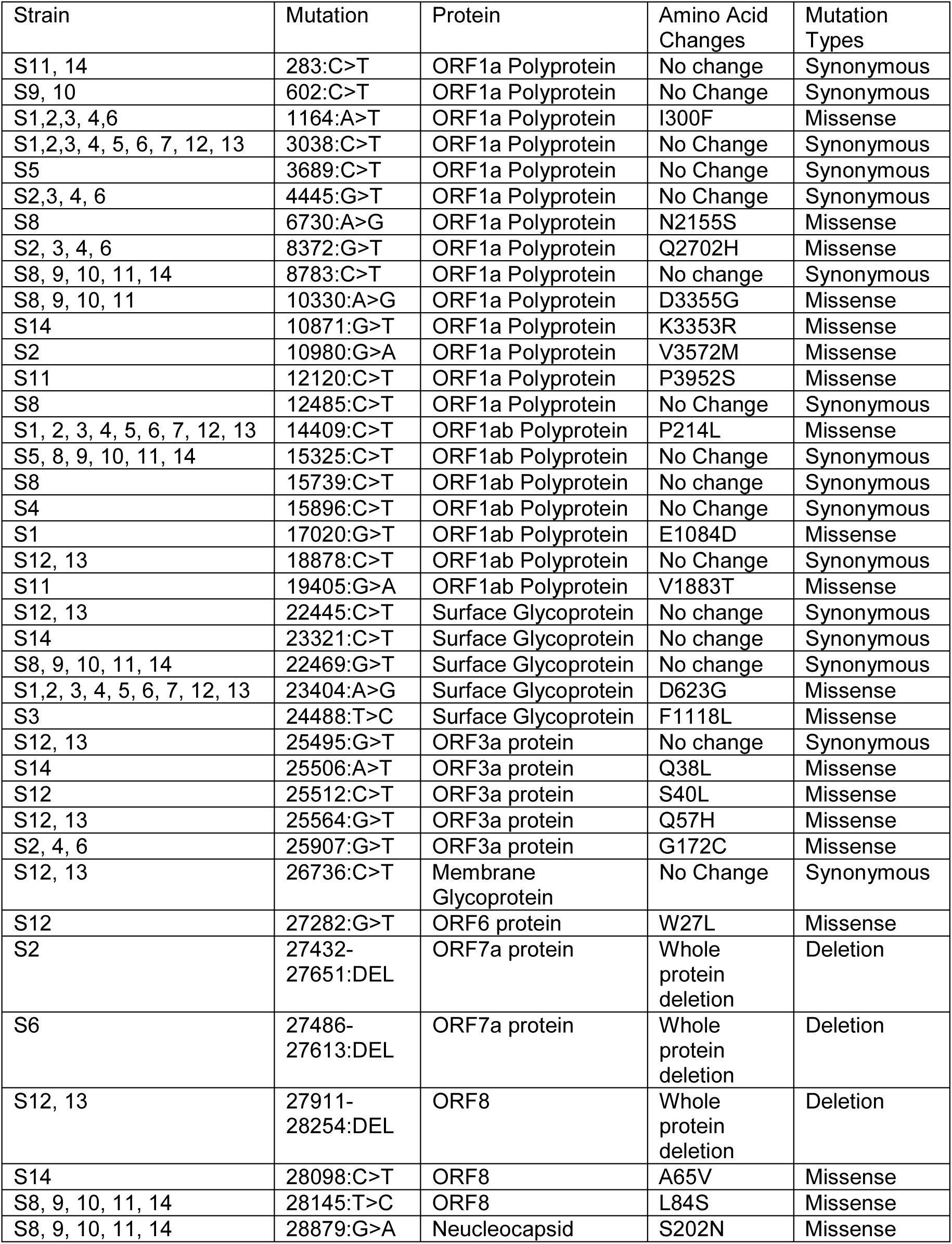
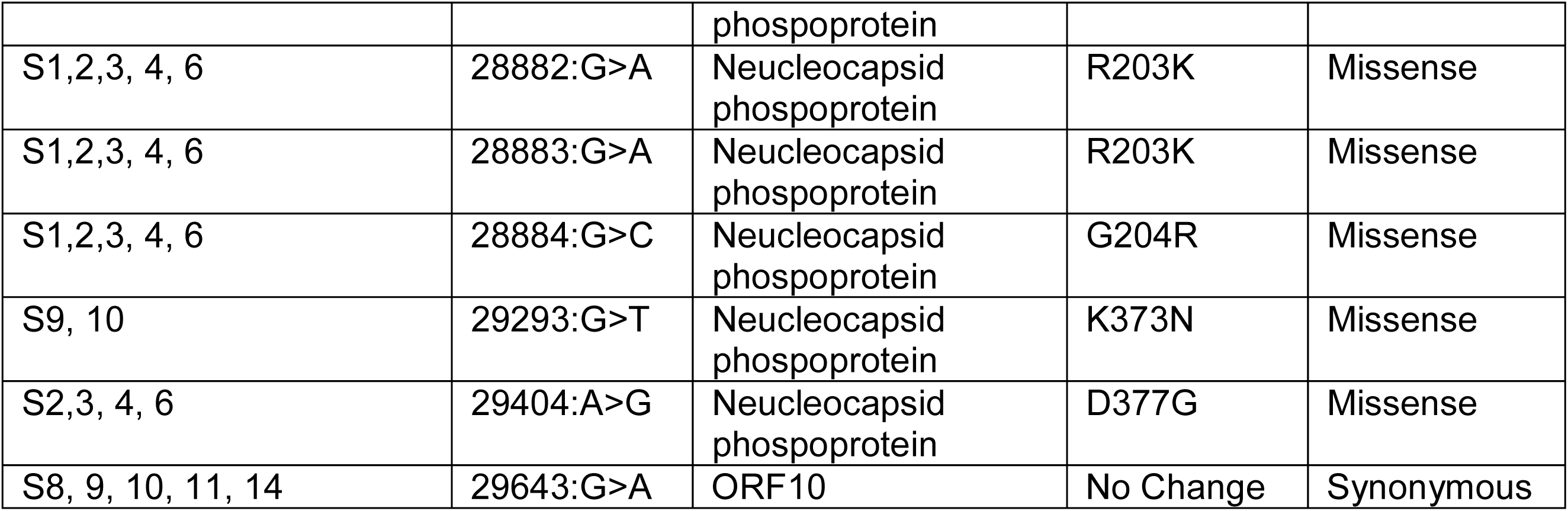
All mutations found in the coding regions of the 14 isolates compared to the reference strain. (Legends: S1: EPI_ISL_437912; S2: EPI_ISL_445213; S3: EPI_ISL_445214; S4: EPI_ISL_445215; S5: EPI_ISL_445216; S6: EPI_ISL_445217; S7: EPI_ISL_445244; S8: EPI_ISL_450339; S9: EPI_ISL_450340; S10: EPI_ISL_4503441; S11: EPI_ISL_450342; S12: EPI_ISL_450343; S13: EPI_ISL_450344; S14: EPI_ISL_450345)

**Fig 2:**
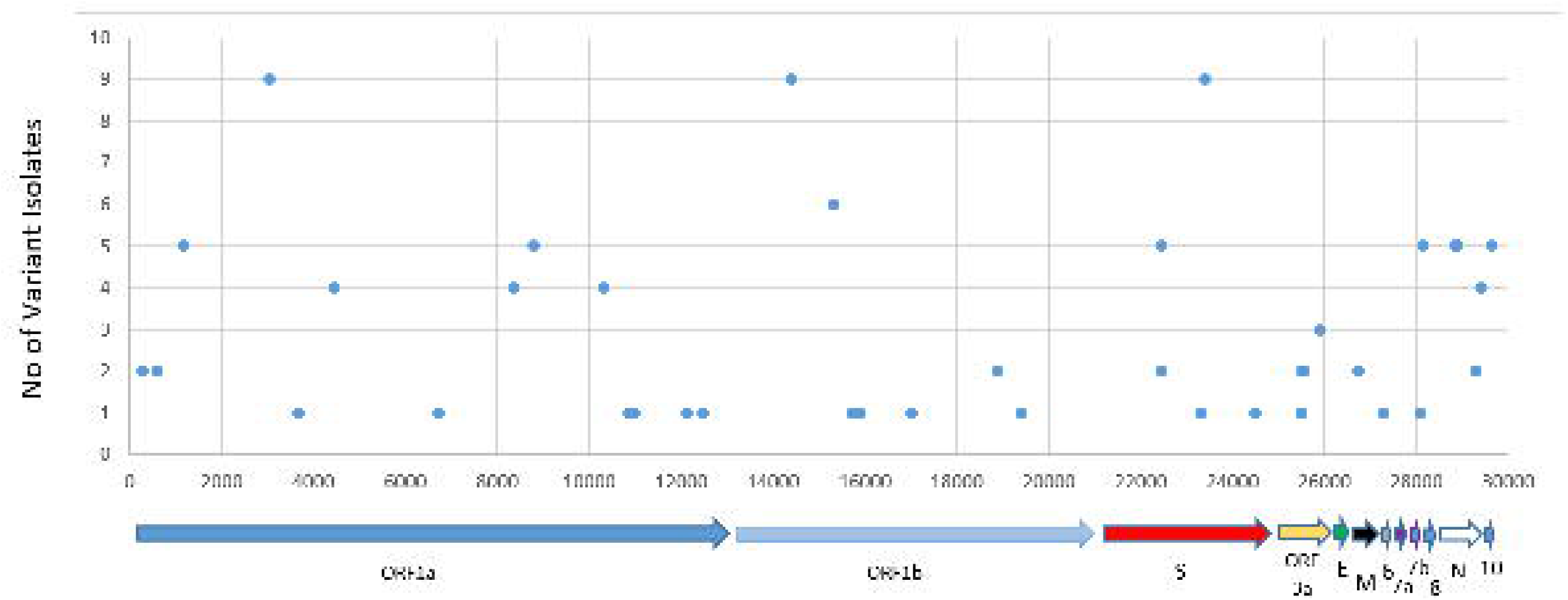
Variations Plot of SARS-CoV-2 in Bangladeshi isolates

### Mutational Effects

Mutational effects analysis of the 24 missense mutations found that 18 mutations were responsible for decreasing structural stability. Mutations located in the ORF1a polyprotein and surface glycoprotein were predicted to decrease the structural stability of both proteins (Table 3). Additionally, three mutations occurring in surface glycoprotein, ORF3a and ORF6 were predicted to alter the molecular consequences, including loss of sulfation in surface glycoprotein and loss of proteolytic cleavage in ORF3a and loss of allosteric site in ORF6 (Table 4 and Supplementary Table 1).

**Table 3:** Prediction of the mutational effects on the structural stability.

| Protein | Amino Acid Changes | SVM2 Prediction Effect | DDG Value (kcal/mol) |
| --- | --- | --- | --- |
| ORF1a Polyprotein | I300F | Decrease | -1.79 |
| ORF1a Polyprotein | N2155S | Decrease | -0.60 |
| ORF1a Polyprotein | Q2702H | Decrease | -0.68 |
| ORF1a Polyprotein | D3355G | Decrease | -0.95 |
| ORF1a Polyprotein | K3353R | Increase | -0.13 |
| ORF1a Polyprotein | V3572M | Decrease | -0.88 |
| ORF1a Polyprotein | P3952S | Decrease | -1.21 |
| ORF1b Polyprotein | P214L | Decrease | -0.83 |
| ORF1b Polyprotein | E1084D | Decrease | -0.75 |
| ORF1b Polyprotein | V1883T | Decrease | -1.46 |
| Surface Glycoprotein | D623G | Decrease | -0.93 |
| Surface Glycoprotein | F1118L | Decrease | -0.81 |
| ORF3a protein | Q38L | Increase | 0.12 |
| ORF3a protein | S40L | Increase | 0.40 |
| ORF3a protein | Q57H | Decrease | -0.90 |
| ORF3a protein | G172C | Decrease | -0.83 |
| ORF6 protein | W27L | Decrease | -0.96 |
| ORF8 | A65V | Increase | 0.02 |
| ORF8 | L84S | Decrease | -2.29 |
| Neucleocapsid phosphoprotein | S202N | Increase | -0.78 |
| Neucleocapsid phosphoprotein | R203K | Decrease | -0.93 |
| Neucleocapsid phosphoprotein | G204R | Decrease | -0.52 |
| Neucleocapsid phosphoprotein | K373N | Increase | -0.10 |
| Neucleocapsid phosphoprotein | D377G | Decrease | -0.44 |

**Table 4:** Prediction of the effects of the mutation on the molecular consequences.

| Protein Name | Mutation | Effects |
| --- | --- | --- |
| Surface Glycoprotein | F1118L | Altered Ordered interface |
|  |  | Altered Disordered interface |
|  |  | Altered DNA binding |
|  |  | Loss of Sulfation at Y1119 |
|  |  | Altered Metal binding |
| ORF3a | G172C | Loss of O-linked glycosylation at S171 |
|  |  | Gain of Disulfide linkage at G172 |
|  |  | Loss of Intrinsic disorder |
|  |  | Altered Transmembrane protein |
|  |  | Altered Ordered interface |
|  |  | Gain of Loop |
|  |  | Loss of Proteolytic cleavage at D173 |
| ORF6 | W27L | Altered Ordered interface |
|  |  | Altered Disordered interface |
|  |  | Loss of Strand |
|  |  | Gain of Helix |
|  |  | Loss of Allosteric site at F22 |
|  |  | Gain of Sulfation at Y31 |
|  |  | Altered DNA binding |
|  |  | Altered Transmembrane protein |

### Prediction and Validation of the Homology Models

In total, three models were generated using the template PDB ID: 6VSB; one model for the spike protein of reference strain, and the two others were for two different mutant isolates from Bangladesh (Fig 3). Two types of mutations were found in the spike proteins of all Bangladeshi isolates, where most of the isolates were found to contain a substitution of D623G. Only one strain, EPI_ISL_445214, found to have two substitutions; one was similar to the previous substitution, and the other was F1118L. The validation assessment scores of these three models were mostly similar to the template, which provided the reliability of these models (Table 5).

**Table 5:** Model Validation assessment score

| Structures | Rampage Score |  | ERRAT Score |
| --- | --- | --- | --- |
|  | Favoured Region | Allowed Region |  |
| Template | 95.8% | 4.1% | 76% |
| China (Ref) | 92.9% | 5.7% | 83% |
| Mutant Model 1 | 92.6% | 5.3% | 84.69% |
| Mutant Model 2 | 92.8% | 5.3% | 83.78% |

**Fig 3:**
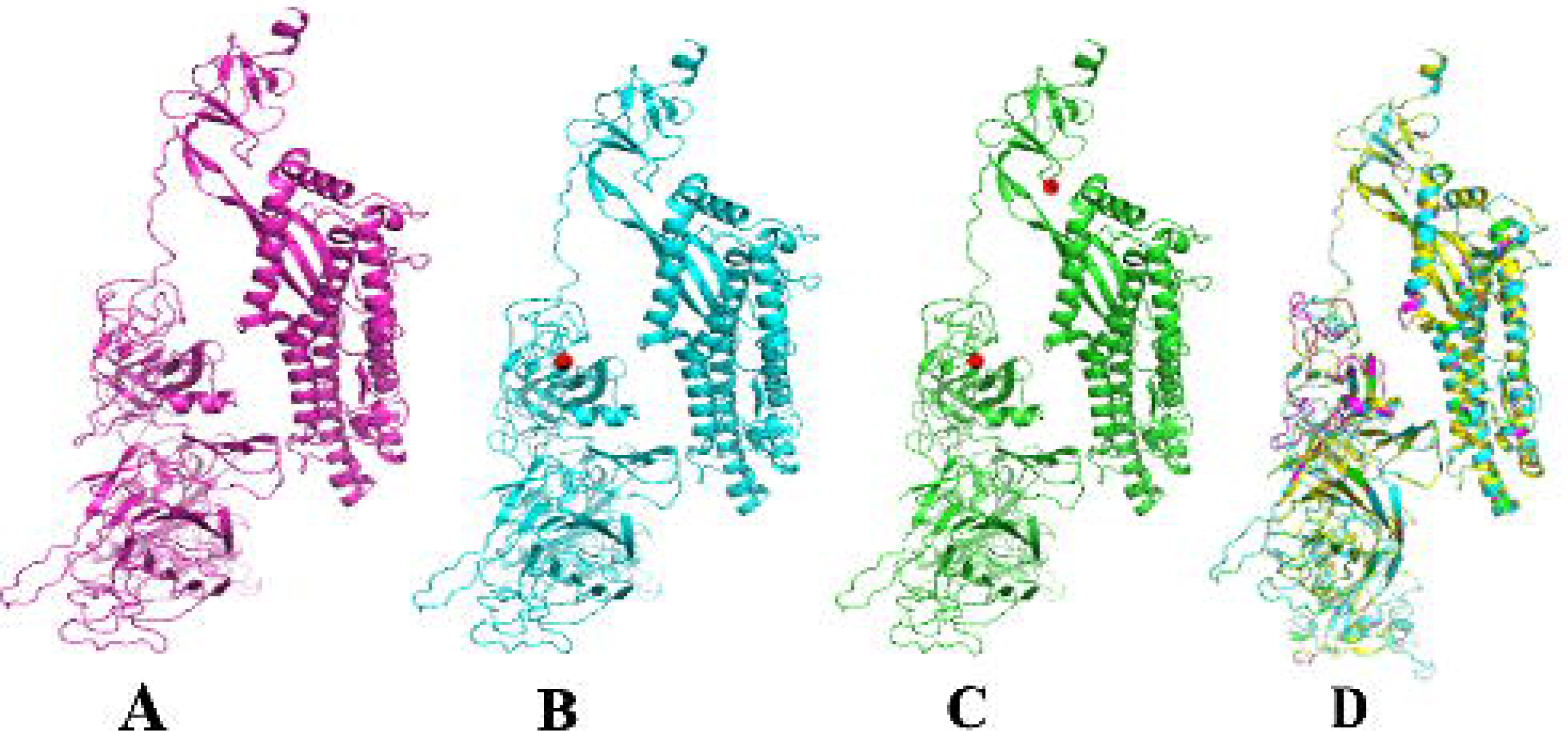
Homology model of the spike proteins; (A) China (B) Model with one mutation: D623G (C) Model with two mutations: D623G and F1118L (D) Superimpose of all model. Here, B and C, red dot represent the mutation site. In D, purple color represents China; the cyan represents a model with one mutation, and the green represents a model with two mutations.

### Analysis of the Interaction Between Spike Proteins and Human ACE2 Receptor

HDOCK server was used to predict the interaction between the above-mentioned 3D models of reference spike proteins along with mutant models and the human ACE2 receptor. Interestingly, this molecular docking analysis revealed that the docking score for the three models against the human ACE2 receptor was similar, and it was −244.42; mutation in the spike proteins do not hamper binding with ACE2 receptor. For three spike protein models, this study found that a domain of spike protein instead of whole protein, amino acid ranging from 345 to 527, was involved in the interactions. This domain was conserved in all isolates resulting in similar interactions with ACE2 (Fig 4).

**Fig 4:**
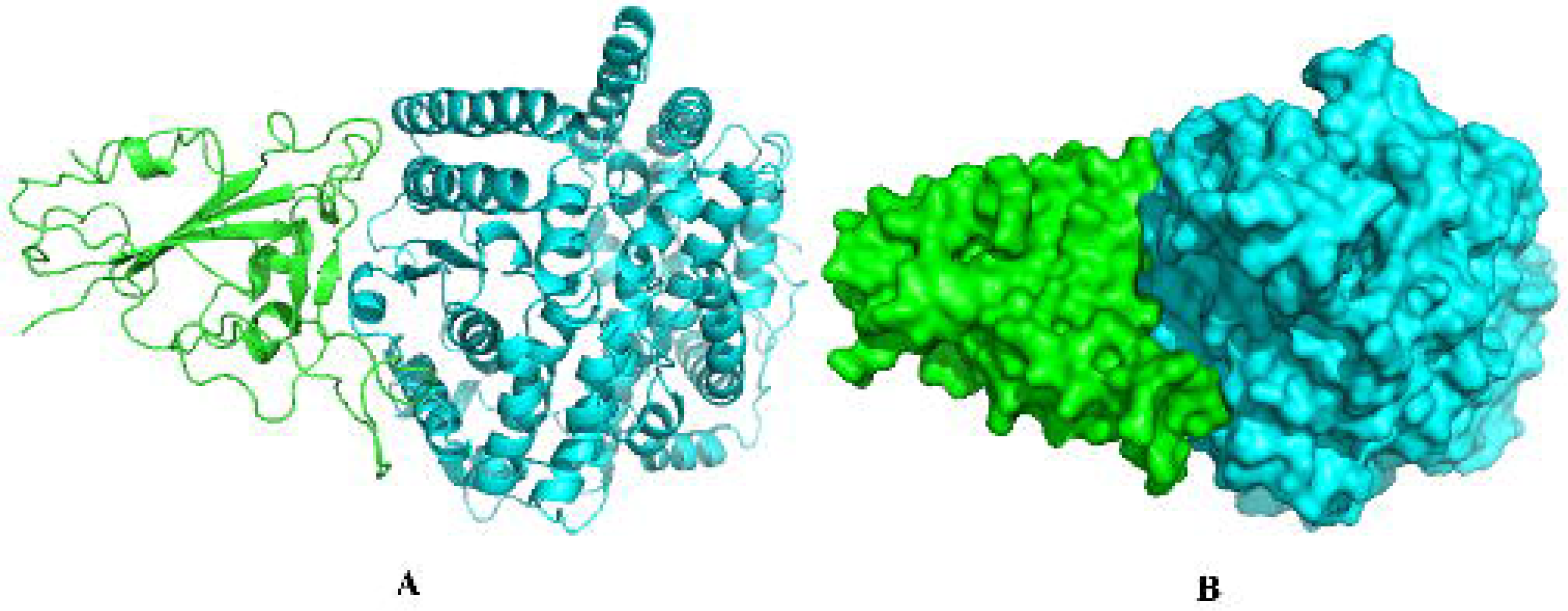
Interaction of Spike protein with ACE2: (A) carton model and (B) Surface model. Here, green represents the binding domain of spike protein, and cyan represents human ACE2.

## Discussion

COVID-19 has become a global challenge for the scientific communities affecting millions of people and taking thousands of lives every day. Scientists worldwide are working hard to combat against SARS-CoV-2, but no significant outcome is obtained [27, 28]. Along with other studies, genetic studies can give a significant clue to understanding the pathogenesis of COVID-19. Together with the critical therapeutic target, the genomic sequence data may provide insights into the pattern of global spread, the diversity during the epidemics, and the dynamics of evolutions, which are crucial to unwind the molecular mechanism of COVID-19 [29]. This study gives insights into the transmission of SARS-CoV-2, genetic diversity of the isolates, and predicts the impacts of mutations in Bangladesh.

It has been reported that, during the COVID-19 outbreak about 600000 people had entered into Bangladesh from the other coutries including Italy and Sapin [30]. The phylogenetic study revealed that the Bangladeshi isolates found in Dhaka were descendent from Europe, and most of the isolates from Chittagong are descendent from the Middle East. However, two isolates of Chittagong were close to the isolates from Dhaka. Dhaka is the capital city of Bangladesh and the sixth most densely populated city in the world. This virus may spread to other regions of the country from this city as it is the central hub of Bangladesh for financial, political, entertainment, and education. The SARS-CoV-2 isolates collected from Chittagong are close the the isolates from the Middle East is not surprising. As most of the migrants from Bangladesh live in Middle East are from Chittagong, and during the COVID-19 outbreak thousands of them returned to their home city [31, 32].

Mutation in the viral genome is a ubiquitous phenomenon for the viruses to escape the host defense. But the mutation rate in SARS-CoV-2 much lower than the other RNA viruses, including seasonal flu viruses [33]. In this study, there was found some variations in the SARS-CoV-2 isolated in Bangladesh, which may affect the epidemiology and pathogenicity of the virus. A total of 42 mutations were identified with a large deletion in the coding regions, where about half were synonymous. Even some isolates found not to encode one or more accessory proteins such as ORF7a, ORF8, and ORF10 caused by a large deletion in the genome. Absent of these accessory proteins may have adverse effects on the viral replication or pathogenesis and the expression of structural protein E [34].

Moreover, ORF8 is involved in the crucial adaptation pathways of coronavirus from human-to-human. At the same time, ORF7a contributes to the viral pathogenesis in the host by inhibiting Bone marrow stromal antigen 2 (BST-2), which restricts the release of coronaviruses from affected cells. Loss of ORF7a causes a much more significant restriction of the virus’s spreading into the host [35, 36]. Loss of these accessory proteins may lead to the virus being less pathogenic, resulting in a meager infection rate and mortality compared to the other countries [34].

Additionally, many variations in structural and non-structural proteins caused substitutions of one or more amino acids were found in the isolates of Bangladesh compared to the reference. Most of the mutations found to affect the structural stability of the proteins rather than alter the molecular functions. Among the structural proteins, most variations were found in Surface glycoproteins (spike) and Nucleocapsid phosphoprotein. Spike proteins play a crucial role in the viral entry into the cell by interacting with the human ACE2 receptor. At the same time, Nucleocapsid phosphoprotein is essential for the packaging of viral genomes into a helical ribonucleocapsid (RNP) and fundamental for viral self-assembly [37, 38]. These functions may not affect much by those mutations, as Mutpred2 predicted that these mutations did not alter any molecular consequences of the proteins.

Moreover, molecular docking analysis revealed that mutations in spike proteins do not affect the interaction with the ACE2 receptor; give us a notion that mutation in the spike protein maybe for the better adaption of the SARS-CoV-2. Thus, therapeutics targeted against the spike protein of SARS-CoV-2 may not give the expected result. This study also identified a domain in the spike protein (amino acid ranging from 345 to 527) involved with human ACE2 receptor interaction rather than the whole protein. This domain was conserved in all isolates reported in Bangladesh, resulting in no effect of the mutations. A recent study identified the receptor-binding domain of spike protein, amino acid ranging from 319 to 541, to interact with the ACE2 receptor, which is similar to our findings [39].

## Conclusion

SARS-CoV-2 isolates from Dhaka and Chittagong were close to European and Mideast lineage. A large deletion in the EPI_ISL_445213, EPI_ISL_445214, EPI_ISL_450343, and EPI_ISL_450344 isolates may explain the less pathogenic result of COVID-19 compared to other countries. Mutations in the spike protein of SARS-CoV-2 may induce more adaptation of this fetal virus; can cause less effective therapeutics if targeted. Our study gives novel insights to understand the SARS-CoV-2 epidemiology in Bangladesh.

## Ethical approval

Not required.

## Data availability

All data supporting the findings of this study are available within the article and its supplementary materials.

## Author Contribution

MH conceived the study. MP and SA designed the study and analyzed the data. MP, MR, MM, and DR performed the experiments. MP wrote the first draft of the manuscript. MH, MP, and SA contributed to the final version of the manuscript. All authors approved the final manuscript.

## Supplementary Files

**Supplementary Table 1:** Mutpred score for all mutations. Scores of < 0.5 indicate no effect on molecular consequences.

