## Supplementary Table 1 for "Genetic analysis of SARS-CoV-2 isolates collected from Bangladesh: insights into the origin, mutation spectrum, and possible pathomechanism"

| Protein | Amino Acid Changes | Mutpred2 Scores |
| --- | --- | --- |
| ORF1a Polyprotein | I300F | .490 |
| ORF1a Polyprotein | N2155S | .460 |
| ORF1a Polyprotein | Q2702H | .387 |
| ORF1a Polyprotein | D3355G | .470 |
| ORF1a Polyprotein | K3353R | .212 |
| ORF1a Polyprotein | V3572M | .185 |
| ORF1a Polyprotein | P3952S | .278 |
| ORF1ab Polyprotein | P214L | .371 |
| ORF1ab Polyprotein | E1084D | .136 |
| ORF1ab Polyprotein | V1883T | .344 |
| Surface Glycoprotein | D623G | .483 |
| Surface Glycoprotein | F1118L | .793 |
| ORF3a protein | Q38L | .306 |
| ORF3a protein | S40L | .213 |
| ORF3a protein | Q57H | .248 |
| ORF3a protein | G172C | .539 |
| ORF6 protein | W27L | .703 |
| ORF8 | A65V | .069 |
| ORF8 | L84S | .311 |
| Neucleocapsid phospoprotein | S202N | .098 |
| Neucleocapsid phospoprotein | R203K | .082 |
| Neucleocapsid phospoprotein | G204R | .158 |
| Neucleocapsid phospoprotein | K373N | .103 |
| Neucleocapsid phospoprotein | D377G | .045 |

**Supplementary Table 1:** Mutpred score for all mutations. Score less than .5 indicates no effect on molecular consequences.
